# *Malpighamoeba* infection compromises fluid secretion and P-glycoprotein detoxification in Malpighian tubules

**DOI:** 10.1101/649483

**Authors:** Marta Rossi, Swidbert R. Ott, Jeremy E. Niven

## Abstract

Malpighian tubules, analogous to vertebrate nephrons, play a key role in insect osmoregulation and detoxification. Tubules can become infected with a protozoan, *Malpighamoeba,* which damages their epithelial cells, potentially compromising their function. Here we used a modified Ramsay assay to quantify the impact of *Malpighamoeba* infection on fluid secretion and P-glycoprotein-dependent detoxification by desert locust Malpighian tubules. Infected tubules have a greater surface area and a higher fluid secretion rate than uninfected tubules. Infection also impairs P-glycoprotein-dependent detoxification by reducing the net rhodamine extrusion per surface area. However, due to the increased surface area and fluid secretion rate, infected tubules have similar total net extrusion per tubule to uninfected tubules. Increased fluid secretion rate of infected tubules likely exposes locusts to greater water stress and increased energy costs. Coupled with reduced efficiency of P-glycoprotein detoxification per surface area, *Malpighamoeba* infection is likely to reduce insect survival in natural environments.

## Introduction

Osmoregulation and excretion are essential for the survival of animals through the homeostatic maintenance of the internal environment and the removal of harmful toxins that impair cellular processes. In insects the Malpighian tubules, which are analogous to the nephrons of the vertebrate kidney, play an important role in osmoregulation as well as in the removal of metabolic waste and xenobiotic substances^1^. Consequently, compromised tubule function may disrupt osmoregulation and impair detoxification, preventing insects from maintaining their internal osmotic environment, and exposing their cells to the waste products of metabolism and to environmental toxins. The impact of such impairment may be particularly severe in those insects that are exposed to environmental toxins as a consequence of feeding directly on plant tissues or their products (such as nectar or pollen) because of the presence of secondary metabolites^2^. Moreover, feeding on plant tissues or their products can expose insects to xenobiotics such as insecticides, fungicides or antibiotics that may act as toxins^3^. Consequently, compromised tubule function may prevent extrusion of these xenobiotics, increasing their effects on insects.

Insect pathogens can interact with xenobiotics such as pesticides affecting survival (reviewed in^3^), though other interactions are known including ingestion of toxins for self-medication^4^,^5^. Interactions between pathogens and xenobiotics, which have been studied primarily in honeybees and bumblebees^4,5^, suggest that pesticide exposure can exacerbate the effects of pathogens. Moreover, pathogens may themselves exacerbate the impacts of xenobiotics, possibly due to trade-offs between the activity of the immune system and detoxification pathways^8^. In most cases, the mechanisms that are affected by Malpighian tubule pathogens and that lead insects to become more susceptible to xenobiotics remain unknown, even in the case of diseases of insect pollinators that provide commercially-important ecosystem services (reviewed in^3^). Consequently, identifying mechanisms by which pathogens could affect removal of xenobiotic compounds is important for understanding insect immune responses, detoxification, Malpighian tubules’ health and overall insects’ health.

In insects, including grasshoppers and honey bees, the Malpighian tubules can become infected by the protozoan *Malpighamoeba*^9^–^14^. This protozoan develops and multiplies primarily in the lumen of the Malpighian tubules before cysts pass into the gut and spread to uninfected individuals through faeces and food contamination^9,13^. Infected tubules appear swollen and cloudy, and their lumen is packed with cysts^9,12^. As the disease progresses, tubule diameter increases in conjunction with the thinning of the epithelium and the destruction of the brush border of the epithelial cells^9,14^. Yet, despite the initial characterisation of *Malpighamoeba* infection over 80 years ago^9,10^ and descriptions of the disease progression^11,12,14^, the effect upon the physiology of Malpighian tubules remains unknown.

Here we compare Malpighian tubule performance of gregarious desert locusts infected with *Malpighamoeba locustae* with that of their uninfected counterparts. We focussed on fluid secretion as well as the removal of hydrophobic organic cations from haemolymph because this requires active transport into the tubule lumen by various types of carriers belonging to the ABC transporter family, including P-glycoprotein^15^. The presence of P-glycoprotein on Malpighian tubules has been demonstrated in several insects, including the desert locust^16–19^. P-glycoproteins are xenobiotic transporters expressed by cells in many tissues throughout the animal kingdom^20^ that transport a broad range of compounds, such as organic cations bigger than 500 Da, and hydrophobic and moderately hydrophobic substances (e.g. alkaloids and quinones)^15^. The function of P-glycoproteins is primarily to reduce the exposure to cells and tissues to harmful compounds. In the case of Malpighian tubules, P-glycoproteins contribute to the excretion of ingested toxic substances^16^.

Here we use the P-glycoprotein substrate rhodamine B^21^ as a proxy for a xenobiotic, to quantify P-glycoprotein activity in Malpighian tubules of the desert locust^19^. P-glycoproteins are likely expressed in the brush border on the apical side of the tubule epithelium^22^, consequently, the destruction of the brush border caused by *Malpighamoeba* infection may compromise xenobiotic substance removal. The infection may also compromise fluid secretion by the Malpighian tubules because damage to the primary and secondary active transporters that move K^+^, Na^+^ and Cl^−^ ions into the lumen that create an osmotic gradient driving the water into the tubule^23^, may be compromised by the destruction of the brush border^9^. To test these hypotheses, we used a modified Ramsay assay^19^ to assess the performance of locust Malpighian tubules.

## Materials and methods

### Animals

Adult desert locusts (*Schistocerca gregaria*, Forsskål, 1775) were taken from a crowded colony maintained at the University of Leicester or were purchased from Peregrine Livefoods (Essex, UK). Locusts were fed with organic lettuce, fresh wheat seedlings and wheat germ *ad libitum.*

### Malpighian tubules, saline and Ramsay assay

Malpighian tubules were removed by cutting the proximal end at ~5 mm from the gut and moved immediately into a 30 μL drop of saline (for the saline composition see^19^) on a 5 cm Sylgard^®^ coated Petri dish, covered with paraffin oil, and fixed with steel pins^19^ (Fig. 1A-D). Isolated tubules ensure that any observed differences in function can be attributed to a direct effect of the infection on the tubules themselves, rather than indirect effects acting at higher levels of organisation^24–26^ e.g. at the level of hormonal control. Each tubule was punctured near the proximal end to allow fluid secretion, and it was allowed to equilibrate for 30 minutes. Subsequently, tubules were incubated in saline containing 60 μM of the P-glycoprotein substrate rhodamine B, and secreted droplets were removed every 30 minutes using a 10 μL pipettor. To ensure that the content of the tubules was fully flushed, we discarded the droplets secreted after 30 and 60 minutes, analysing only the droplets secreted at 90 minutes. These droplets were photographed with a digital camera (Canon EOS 7D; Canon, Tokyo, Japan) mounted on a stereoscopic microscope (Nikon SMZ-U; Nikon Corp., Tokyo, Japan) (Fig. 1E). Photobleaching of rhodamine was prevented using a custom dark box. Using the image processing program ImageJ v.1.51p^27^, we measured the volume of the droplets secreted to calculate the fluid secretion rate (nL/min) by dividing the droplet volume by the time of secretion (30 min), and we estimated the rhodamine concentration (μM) of the droplets from their colour intensity using a known calibration curve^19^. We then calculated the net rhodamine extrusion rate (fmol/min) by multiplying the secretion rate by the rhodamine concentration to estimate the moles of rhodamine extruded per minute.

**Figure 1.**
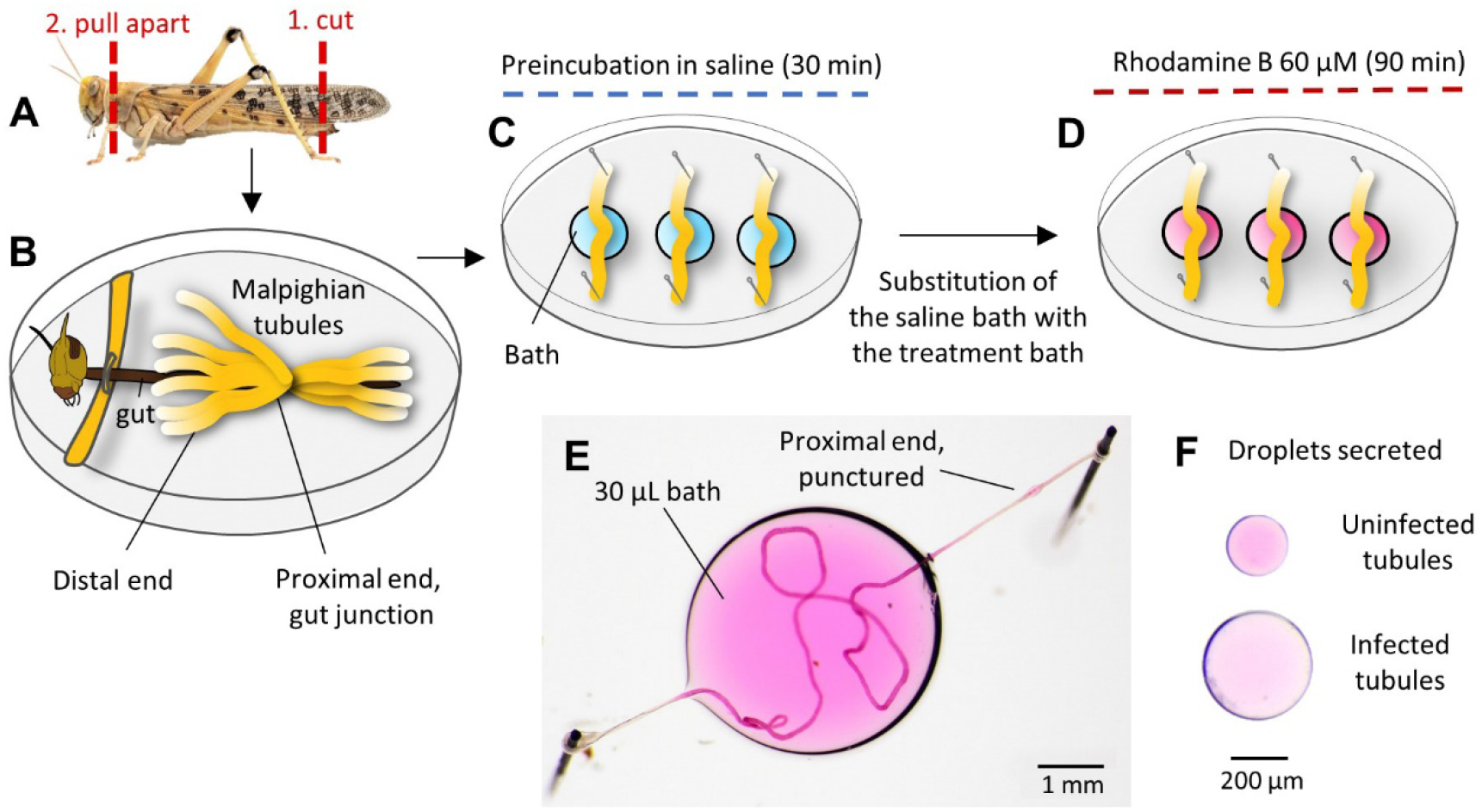
Rhodamine B extrusion assay of Malpighian tubule function. **A)** Locust dissection. **B)** Locust dissected with Malpighian tubules exposed. **C)** Three tubules removed from each locust and placed in a saline bath. The proximal end of each tubule was punctured with a sharpened capillary tube to allow the luminal fluid to be secreted. **D)** After 30 minutes of preincubation, the saline was removed from the bath wells and replaced with 60 μM rhodamine B in saline; we discarded the droplets secreted after 30 and 60 minutes, analysing only those secreted at 90 minutes. **E)** An example of a tubule incubated in the treatment bath containing rhodamine. The proximal end of the tubule is punctured to allow fluid secretion. **F)** Examples of droplets secreted after 90 minutes of incubation by uninfected (above) and infected (below) Malpighian tubules. These droplets were removed from the puncture site in the proximal end and then placed onto a Petri dish for photography.

At the end of the assay, we measured the tubule diameter and length, and the length of tubule in contact with the treatment bath (Table 1). From these measurements, we calculated the surface area of the tubule and the surface area in contact with the treatment bath. We removed three Malpighian tubules from eight uninfected locusts and nine locusts infected with *Malpighamoeba locustae* (Amoebidae, Sarcodina). One infected tubule was damaged during the experiment and therefore excluded from further analysis. This produced a total of 24 uninfected and 26 infected tubules. Each removed tubule was incubated in rhodamine B individually. We selected tubules with similar level of infection, excluding those that were visibly damaged by globular melanised encapsulations that fuse several tubules together.

**Table 1.** Summary of the statistical models to compare morphological differences between uninfected and infected tubules (health status). Infected tubules are longer and broader with a larger surface area than uninfected tubules. The surface area exposed to the incubation bath was greater in infected tubules than uninfected ones. Each morphological trait response was fitted by a GLMM with gamma error distribution and identity link function. Locust ID was included as random effect. Effect estimates for the independent two-level factor “health status” (uninfected vs. infected) report the estimated difference in the group means of tubules from uninfected and infected individuals, together with the standard error of the difference, the *t*-statistic and *p*-value of the Null Hypothesis that means are the same.

| Morphological traits | Fixed effects | Estimate | s.e. | *t* | *p* |  |
| --- | --- | --- | --- | --- | --- | --- |
| Length (mm) | Intercept (uninfected) | 21.09 | 0.67 | 31.35 |  |  |
|  | Health status | 9.84 | 1.09 | 9.00 | **<0.001** | *** |
| Diameter (µm) | Intercept (uninfected) | 56.36 | 7.36 | 7.66 |  |  |
|  | Health status | 48.88 | 8.81 | 5.55 | **<0.001** | *** |
| Surface (mm<sup>2</sup>) | Intercept (uninfected) | 3.87 | 0.70 | 5.52 |  |  |
|  | Health status | 6.17 | 0.87 | 7.05 | **<0.001** | *** |
| Surface in bath (mm<sup>2</sup>) | Intercept (uninfected) | 2.16 | 0.26 | 8.31 |  |  |
|  | Health status | 4.51 | 0.50 | 9.08 | **<0.001** | *** |

### Statistical Analysis

All statistical analyses were conducted in R version 3.4.1 (R Core Team, 2018). The distributions of the dependent variables analysed had a theoretical lower bound of zero and showed clear heteroscedasticity and positive skewness. To account for these distributional properties, we fitted Generalized Linear Mixed Model (GLMM) with Gamma error distributions and “inverse” or “identity” link functions accordingly to the distribution of the data (`glmer` function, package ‘lme4’^28^). Adequacy of the model fits was assessed from diagnostic plots of the standardised residuals (quantile-quantile and residuals over fitted). We investigated: (i) the effect of surface and infection on the fluid secretion rate; (ii) the effect of fluid secretion rate and infection on rhodamine concentration; and (iii) the effect of fluid secretion rate and infection on the net rhodamine extrusion. The predictors were scaled to standard deviation (SD)=1, and centred on mean=0, while the dependent variables were scaled to SD=1 without re-centring. To quantify these effects, we selected the most parsimonious model from candidate models of different complexity using the Bayesian information criterion (BIC)^29^ (Tables 2–4). To account for the nested structure of data, we included the individual locust as random intercepts in the model.

Effect estimates for the independent two-level factor “health status” (uninfected vs. infected) report the estimated difference in the group means of tubules from uninfected and infected individuals, together with the standard error of the difference, the *t*-statistic and *p*-value of the Null Hypothesis that means are the same. Effect estimates for continuous independent variables (“tubule surface” and “secretion rate”) report the estimated change in the value of the dependent variable, in standard deviation units, when changing the value of the independent variable by one standard deviation.

## Results

We compared the morphology of the Malpighian tubules of uninfected desert locusts with those infected by the protozoan *Malpighamoeba locustae* (Fig. 2). Uninfected Malpighian tubules were transparent with a distinct lumen (Fig. 2A,B), whereas the lumen of infected tubules were filled with *M. locustae* cysts throughout their length and had a thinner and less clearly defined wall than that of uninfected tubules (Fig. 2C,D). The rhythmic movements of the tubules were also slower and less pronounced in infected compared with uninfected locusts (Supplementary Videos S1,2). We measured the length and diameter of Malpighian tubules in both uninfected and infected locusts, and calculated their surface area from their length and diameter (Fig. 3). Infected tubules were significantly longer and broader with a larger outer surface area than uninfected tubules (Table 1; Fig. 3), and as a result a greater surface area was exposed to the incubation bath in infected compared to uninfected tubules (Table 1).

**Figure 2.**
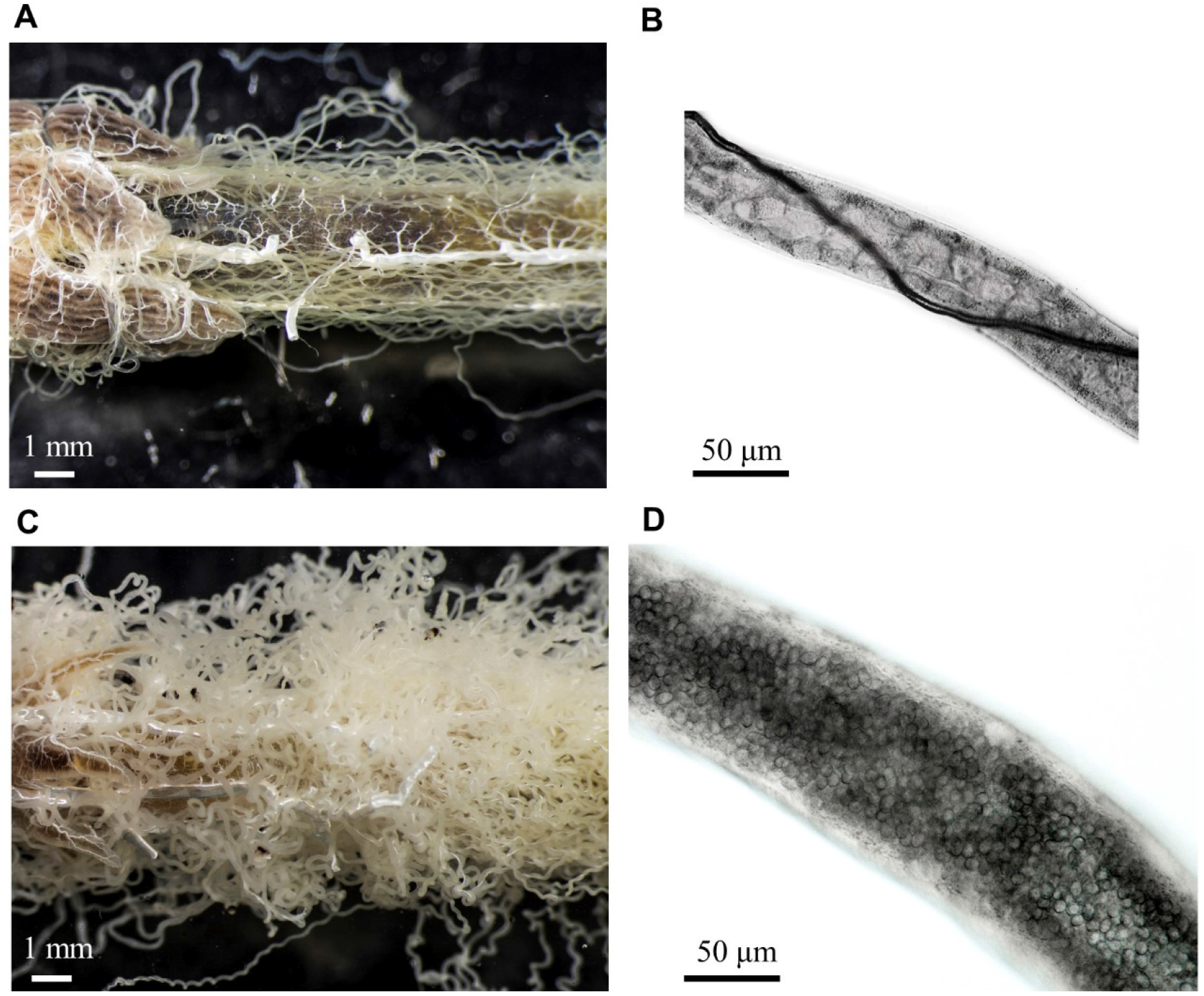
The Malpighian tubules of Malpighamoeba-infected locusts differ from those of uninfected locusts in appearance. (A) Uninfected tubules are thinner and more transparent. (B) The lumen of uninfected tubules is free from cysts. The dark structure running along the length of the tubule in this image is a tracheal branch. (C) Infected tubules are swollen and cloudy. (D) The lumen of infected tubules is filled with cysts.

**Figure 3.**
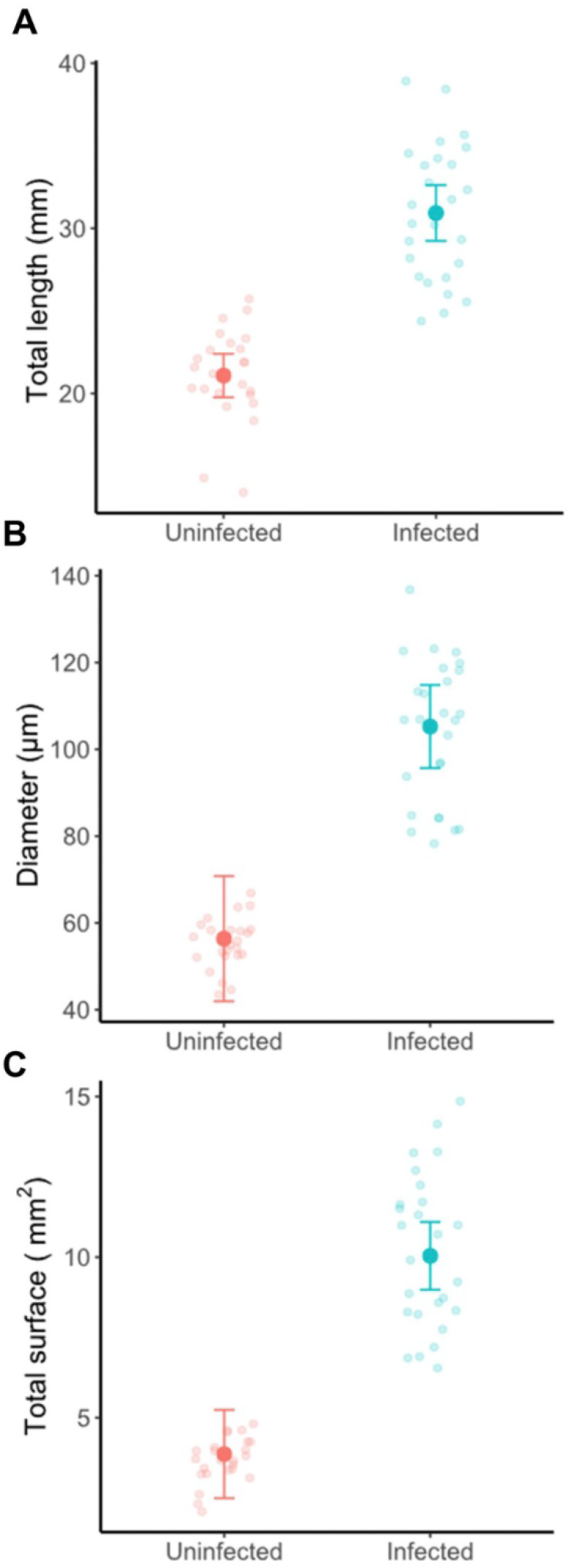
The Malpighian tubules of Malpighamoeba-infected locusts differ from those of uninfected locusts in length, diameter and surface area. Infected locusts had longer (A) and wider (B) tubules than uninfected locusts. (C) Consequently, the surface area of infected tubules was greater than that of uninfected tubules. Large filled circles with error bars represent mean estimates from GLMM fits with the 95% confidence interval. Small circles represent the raw data.

One function of the Malpighian tubules is to maintain osmotic balance through secretion of water and ions during diuresis^23^. To test whether infection with *M. locustae* affects the fluid secretion rate of tubules, we quantified the relationship between fluid secretion rate and surface area for uninfected or infected tubules (Fig. 4). We selected the most parsimonious model from four candidate models with decreasing complexity (Table 2A). We found that surface area positively influences the fluid secretion rate independently of the health status of tubules (Table 2B, Fig. 4). Thus, for a given unit of surface area tubules possess similar secretion rates, and consequently infected locusts with distended tubules tend to secrete more fluid (1.019 nl/min, 95% CI = [0.65, 1.39]) than uninfected locusts with smaller tubules (0.50 nl/min [0.13, 0.88]).

**Figure 4.**
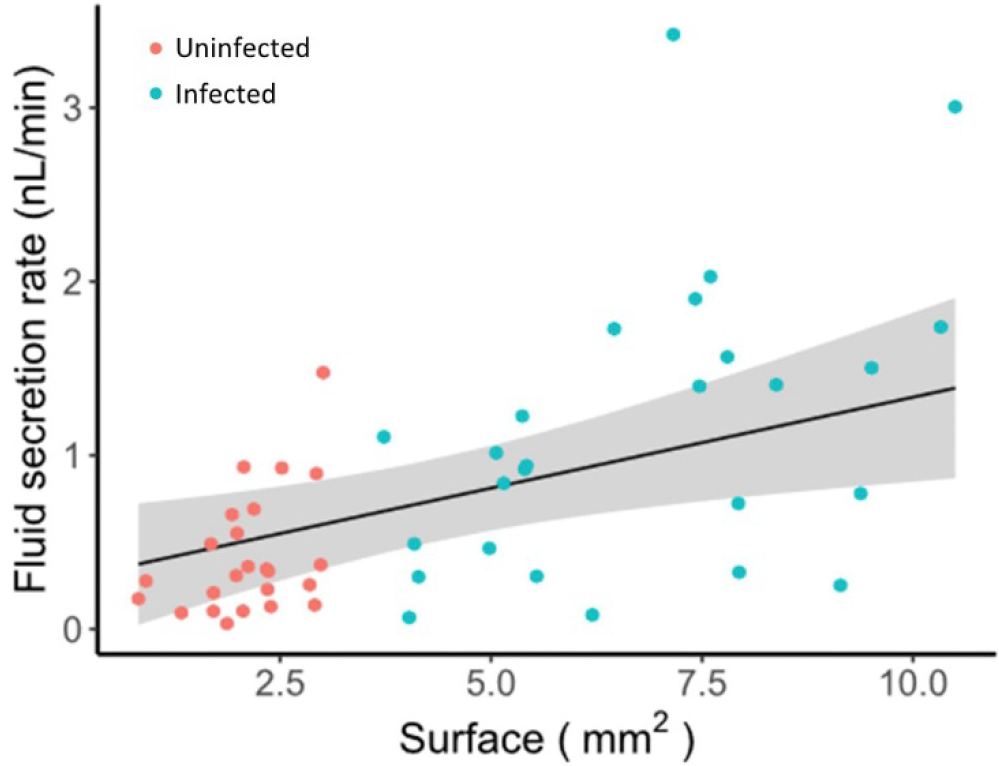
*Malpighamoeba* infection affects the fluid secretion rate of Malpighian tubules. Tubule surface area positively correlates with fluid secretion rate, therefore infected tubules with larger surface tend to secrete more fluid than the smaller uninfected tubules.

**Table 2.** Summary of the statistical models to investigate the effect of surface and health status on the fluid secretion rate. **A**) Comparison of statistical models predicting the fluid secretion rate (SR; scaled to S.D.=1) in relation to tubule surface area (scaled to S.D.=1 and mean centred on 0) and/or health status (uninfected, infected). The SR response was fitted by a GLMM with gamma error distribution and identity link function. Locust ID was included as random effect. According to the BIC, surface area alone explains the observed secretion rate (model in bold). **B**) Summary of the statistical model predicting SR in relation to surface area. Effect estimate for the continuous independent variable “surface” reports the estimated change in the value of the dependent variable, in standard deviation units, when changing the value of the independent variable by one standard deviation.

| A. Candidate models |  | df | BIC |
| --- | --- | --- | --- |
| SR ~ surface * health status |  | 6 | 54.614 |
| SR ~ surface + health status |  | 5 | 56.135 |
| **SR ~ surface** |  | **4** | **52.254** |
| SR ~ health status |  | 4 | 55.046 |

| B. Fixed effects | Estimate | s.e. | t | p |
| --- | --- | --- | --- | --- |
| Intercept | 0.699 | 0.112 | 6.26 |  |
| surface | 0.267 | 0.096 | 2.77 | **0.00566** ** |

In addition to fluid secretion, Malpighian tubules also contribute to the extrusion of metabolic waste and xenobiotics from the haemolymph^23^. The altered morphology (Figs. 2,3) and increased fluid secretion (Fig. 4) may affect the ability of infected Malpighian tubules to extrude xenobiotics. To test this, we quantified the extrusion of the P-glycoprotein substrate rhodamine B^18,19,21^ during tubule incubation. Rhodamine B was transported into the tubule lumen (Fig. 5A), and its concentration in the droplets secreted by tubules was higher in uninfected than infected tubules (uninfected: 211.54 μM [127.35, 624.32]; infected: 77.12 μM [61.97, 102.08]; Table 3, Fig. 5B). The secretion rate had a negligible effect on the rhodamine concentration, so that the lower rhodamine concentration in droplets secreted by the infected tubules cannot be attributed to their higher secretion rates and the consequent dilution (Table 3).

**Figure 5.**
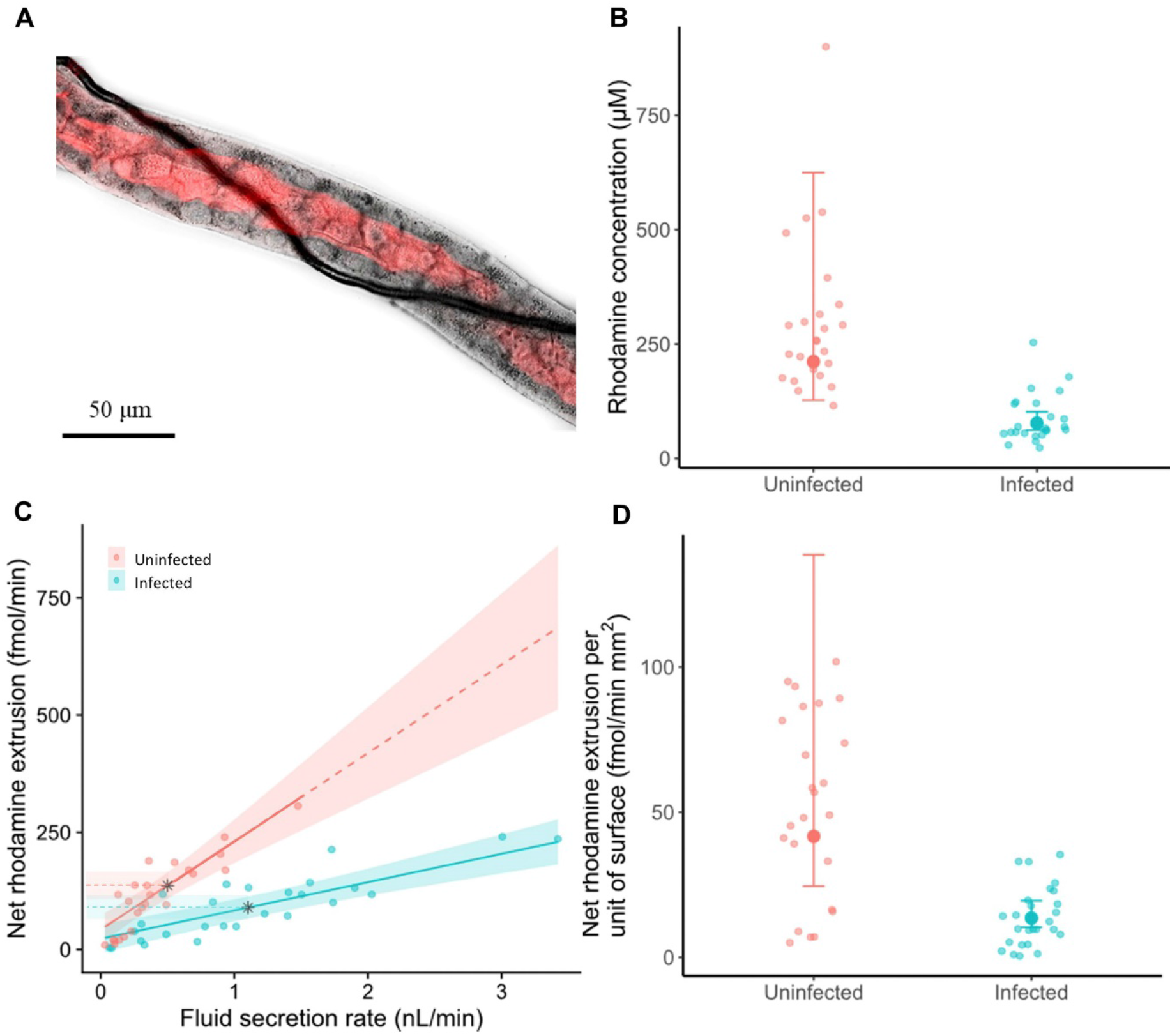
*Malpighamoeba* infection affects rhodamine B extrusion by Malpighian tubules. **A)** The uninfected tubule shown in Fig. 1B transporting rhodamine B. **B**) Uninfected tubules secreted more concentrated droplets than the infected tubules. **C**) The secretion rate positively correlates with the net extrusion rate of rhodamine B, but the steepness is more accentuated in the uninfected tubules compared to the infected ones. Black stars represent the mean fluid secretion rate for uninfected and infected tubules. **D**) The net rhodamine extrusion rate per unit surface area is higher in uninfected than infected tubules. Small circles represent the raw data, large filled circles represent means estimates, and lines represent the regression line fitted from GLMM. Shaded areas and error bars represent the 95% confidence interval. All the values are back transformed to the original scale.

**Table 3.**
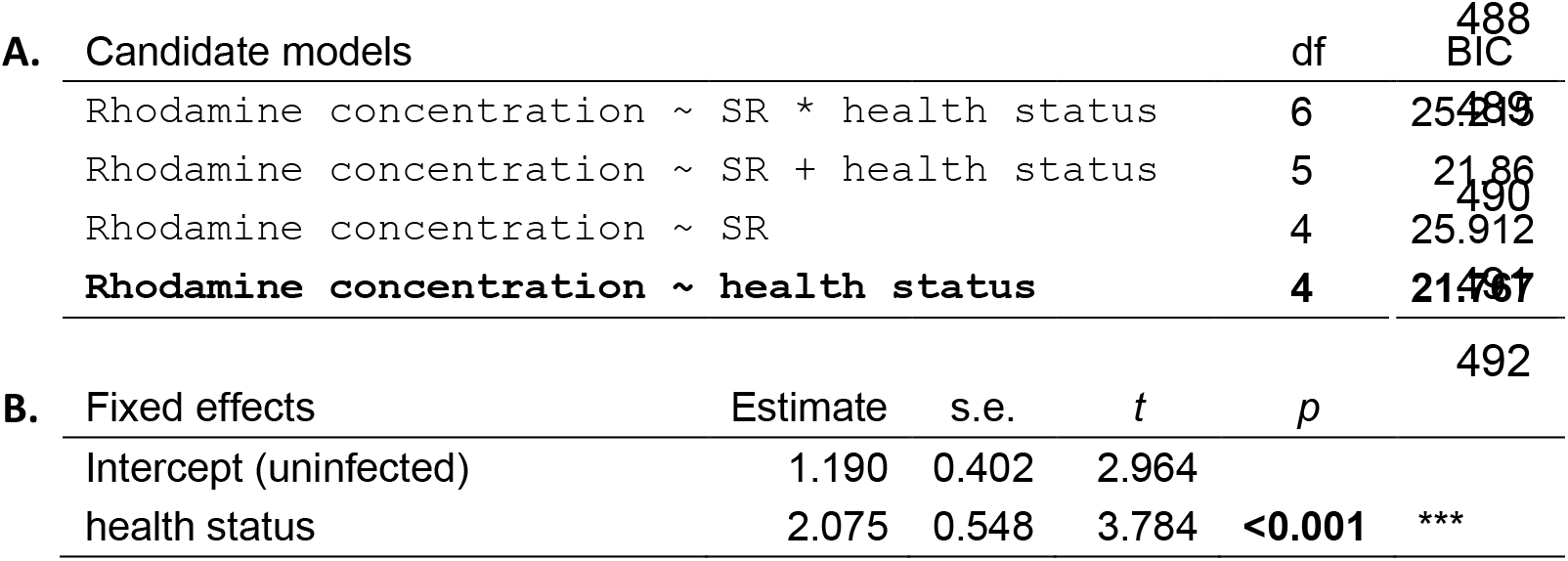
Summary of the statistical models to investigate the effect of fluid secretion rate and health status on the rhodamine concentration of the droplets secreted. **A**) Comparison of statistical models predicting the rhodamine concentration (scaled to S.D.=1) in relation to the fluid secretion rate (SR; scaled to S.D.=1 and mean centred on 0) and/or health status (uninfected, infected). The rhodamine concentration response was fitted by a GLMM with gamma error distribution and inverse link function. Locust ID was included as random effect. According to the BIC, health status alone explains the observed rhodamine concentration (model in bold). **B**) Summary of the statistical model predicting the rhodamine concentration in relation to health status. Effect estimates for the independent two-level factor “health status” (uninfected vs. infected) report the estimated difference in the group means of tubules from uninfected and infected individuals, together with the standard error of the difference, the *t*-statistic and *p*-value of the Null Hypothesis that means are the same.

The net rhodamine extrusion rate (femtomoles of rhodamine extruded per minute) increased with the fluid secretion rate, but this increase was less pronounced in infected tubules than in uninfected ones (Table 4A,B; Fig. 5C). Consequently, for a given fluid secretion rate, infected tubules extrude less rhodamine. At the mean tubule fluid secretion rate of 0.79 nl/min, the mean net extrusion rate of uninfected tubules was 191.29 fmol/min [153.92, 228.65], whereas the mean net extrusion rate of infected tubules was less than half of this (70.75 fmol/min [46.36, 95.15]). However, the tendency of infected tubules to have a greater fluid secretion rate (uninfected, mean: 0.50 nl/min; infected, mean: 1.019 nl/min) counteracts, at least partially, the reduced net extrusion at any given fluid secretion rate. Therefore, the overall rate of rhodamine extruded by a typical infected tubule (89.58 fmol/min [65.92, 113.25] and a typical uninfected tubule (136.67 fmol/min [108.57, 164.77]) are broadly comparable (Table 4A,B; Fig. 5C, black stars). Importantly, because the net rhodamine extruded is a dependent variable of the fluid secretion, the increased fluid secretion rates of infected Malpighian tubules mean that infected and uninfected tubules have comparable overall extrusion rates. The reduction in rhodamine extrusion by infected tubules in comparison to uninfected tubules suggests decreased extrusion per unit surface area. We quantified the net rhodamine extrusion per unit surface area and found that it was indeed significantly reduced in infected tubules (13.53 fmol/min·mm^2^, [10.36, 19.51]) compared with uninfected tubules (41.72 fmol/min·mm^2^ [24.56, 138.64]; *t*=3.97, *p*<0.001; Fig. 5D).

**Table 4.** Summary of the statistical models to investigate the effect of fluid secretion rate and health status on the net rhodamine transport. **A**) Comparison of statistical models predicting the net rhodamine transport (scaled to S.D.=1) in relation to the fluid secretion rate (SR; scaled to S.D.=1 and mean centred on 0) and/or health status (uninfected, infected). The net rhodamine transport response was fitted by a GLMM with gamma error distribution and identity link function. Locust ID was included as random effect. According to the BIC, the interaction between SR and health status best explains the net rhodamine transport (model in bold). **B**) Summary of the statistical model predicting the net rhodamine transport in relation to SR and health status. Effect estimates for the independent two-level factor “health status” (uninfected vs. infected) report the estimated difference in the group means of tubules from uninfected and infected individuals, together with the standard error of the difference, the *t*-statistic and *p*-value of the Null Hypothesis that means are the same. Effect estimate for the continuous independent variable “SR” reports the estimated change in the value of the dependent variable, in standard deviation units, when changing the value of the independent variable by one standard deviation.

| A. Candidate models |  | df | BIC |
| --- | --- | --- | --- |
| **Net rhodamine transport ~ SR \* health status** |  | **6** | **12.943** |
| Net rhodamine transport ~ SR + health status |  | 5 | 28.363 |
| Net rhodamine transport ~ SR |  | 4 | 33.871 |
| Net rhodamine transport ~ health status |  | 4 | 67.018 |

| B. Fixed effects | Estimate | s.e. | *t* | *p* |  |
| --- | --- | --- | --- | --- | --- |
| Intercept (uninfected) | 1.507 | 0.150 | 10.032 |  |  |
| SR | 1.098 | 0.165 | 6.648 | **<0.001** | *** |
| Health status (at mean SR) | -0.950 | 0.179 | -5.316 | **<0.001** | *** |
| SR × health status | -0.746 | 0.175 | -4.269 | **<0.001** | *** |

## Discussion

We determined how infection by the protozoan *Malpighamoeba locustae* affects the performance of insect Malpighian tubules using gregarious desert locusts *(Schistocerca gregaria)* as a model system. We compared the performance of infected and uninfected tubules in terms of secretion and xenobiotic extrusion. By doing so, our results directly quantify the impact of a common pathogen upon major components of insect osmoregulation and xenobiotic excretion, namely the Malpighian tubules and the P-glycoproteins they express. Infected tubules differed from uninfected tubules in morphology being longer, wider, and possessing a greater surface area. The infected tubules also differed in behaviour from uninfected tubules showing less pronounced and slower rhythmic movements (Supplementary Videos S1,2). We assessed two key aspects of the physiological performance of tubules; their fluid secretion and net extrusion of rhodamine B, which is a P-glycoprotein substrate^21^. Our analysis showed that infected tubules with a larger surface area secrete more fluid per unit time. Therefore, as a consequence of their greater surface area, infected tubules tend to secrete more fluid per unit time than uninfected tubules. Despite this increased secretion, however, the net extrusion of rhodamine per unit surface area is far lower in infected compared with uninfected tubules.

Fluid secretion from tubules is powered by the apical V-ATPase, which creates an electrochemical gradient that secondarily drives Na^+^, K^+^, and Cl^−^ ion movement into the lumen, followed by passive water movement^23^. The V-ATPase is expressed within the apical brush border of the tubule epithelium^30,31^. The *Malpighamoeba* trophozoites destroy the apical brush border of Malpighian tubules^9,14^, raising the possibility that the V-ATPase is reduced or absent. Consequently, the greater fluid secretion rate of the infected compared to the uninfected tubules is at first quite surprising. However, the reduction of the brush border and consequent change in the composition of cellular cytoplasm may produce ionic gradients that permit continued transcellular water movement, which may be sufficient to produce similar rates of fluid secretion rate per unit of surface area.

Indeed, the epithelial cell is no longer a sealed compartment, and the cytoplasm is in direct contact with the lumen of the tubule. Therefore, even without the V-ATPase, the ions could move through the Na^+^:K^+^:2Cl^−^ cotransporter from the haemolymph directly into the lumen, and in turn the water may follow osmotically. Alternatively, the thinning of the epithelium or the damage of the brush border in infected tubules may compromise the septate junctions, which exist between epithelial cells^32^, promoting water movement through a paracellular pathway.

Trans-tubular rhodamine transport is predominantly dependent on P-glycoprotein transporters^19,21^, although rhodamine can passively diffuse across membranes^21^, and in human cell lines (Calu-3) it can interact with organic cation transporters such as OCT3 and OCTN1,2^33^. P-glycoproteins are expressed in the brush border of tubule epithelial cells^22^, though they may be present on the basal side^16^. The destruction of the brush border by the *Malpighamoeba* trophozoites^9,14^ likely impairs P-glycoprotein transport. However, the greater fluid secretion rate of infected tubules will decrease their luminal rhodamine B concentration in comparison to uninfected tubules. A lower luminal rhodamine concentration reduces the back diffusion^19,34^ of rhodamine from the lumen to the bath solution, thereby increasing its net rhodamine B extrusion. Nevertheless, infected tubules extrude less rhodamine B for a given rate of fluid secretion and less rhodamine B per unit surface area than do uninfected ones. Infected tubules do extrude rhodamine B, which may be via diffusion down a concentration gradient or by relying on other organic cation transporters, as suggested by studies of human cell lines^33^.

The overall consequence of the reduced net extrusion of rhodamine per unit of surface area coupled with the increased fluid secretion rate is that the total net extrusion of rhodamine per tubule is similar in infected and uninfected tubules. This is consistent with the finding that infection with *Malpighamoeba* does not increase the susceptibility of grasshoppers *(Melanoplus sanguinipes)* to the insecticide cypermethrin, a P-glycoprotein substrate, within one hour of application^35,36^. However, tubules of heavily infested locusts present globular melanised encapsulations that can fuse together and to adjacent tissues forming tissue masses that can no longer contribute to excretion^9,12^ (Fig. 6). Therefore, it is possible that these encapsulations reduce the number of functional tubules, further compromising locust excretion.

**Figure 6.**
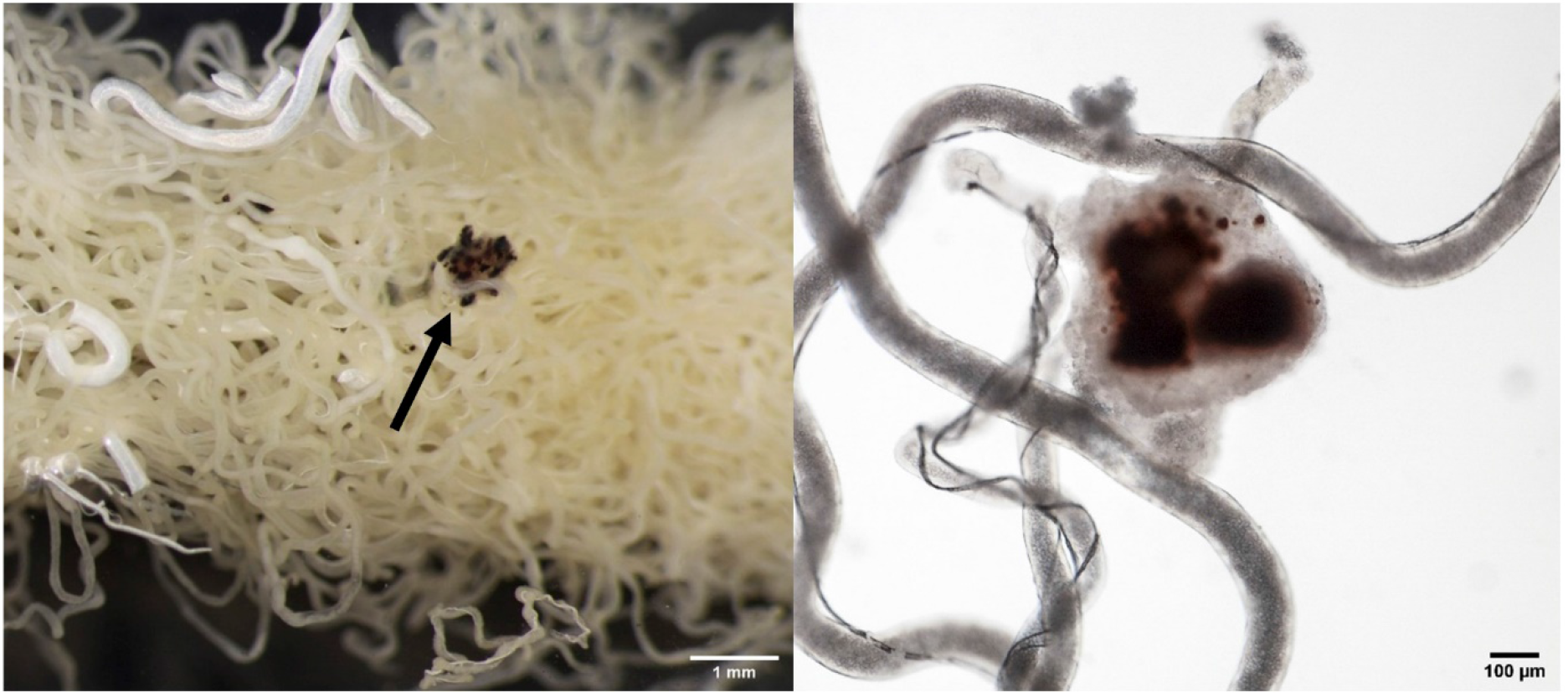
Locusts Malpighian tubules infected with *Malpighamoeba locustae* contain globular melanised encapsulations. **A)** Malpighian tubules fused together with melanised encapsulations indicated by the arrow. B) Isolated encapsulation.

Changes in feeding behaviour may also contribute to the ability of locusts to compensate for *Malpighamoeba* impairment of P-glycoprotein detoxification; infection reduces the amount of food that locusts ingest^9^ (M. Rossi pers. obs.). Reducing food intake may reduce the exposure of infected insects to noxious substances. Locusts infected with *Malpighamoeba* experience higher fluid secretion to achieve similar levels of toxin extrusion as uninfected locusts. Consequently, the reduced food intake may reduce infected locusts’ exposure to noxious substances that their compromised detoxification pathways cannot remove without substantial water loss into the gut. This may be particularly important for generalist herbivores like desert locusts that feed on a broad variety of plant species, many of which contain toxins^37^. Alternatively, reduced food intake in infected locusts may be driven by the activation of the immune system^8^. Both their immune system and their ability to detoxify noxious food substances rely on pathways that involve the antioxidant glutathione^38–40^. Consequently, competition for glutathione between these two physiological pathways could lead to impaired detoxification when the immune system is already activated^8^. Whether due to a direct reduction in feeding or as an indirect consequence of a trade-off with the immune response, reduced food intake lowers the risk of ingesting toxins but also prevents locusts acquiring water and nutrients.

A reduced food intake combined with an increased fluid secretion rate poses infected locusts a difficult challenge. In adaptation to the dry and hot environment that they occupy, desert locusts must balance their water secretion to avoid dehydration^41^. Diuretic and antidiuretic hormones play an important role in the control of the tubular fluid secretion^42^. However, *Malpighamoeba* infection may damage the hormone receptors, decreasing the ability of the insect to respond to the circulating hormones^43^. The primary urine secreted by the Malpighian tubules is isosmotic to the haemolymph, and the composition must be adjusted prior to excretion^23^. Most of the filtered water, ions and metabolites are reabsorbed in the anterior hindgut and in the rectum^23^. The reabsorption of Na^+^, K^+^, Cl^−^ ions and hence water is an active process consuming metabolic energy^41,44^. To prevent desiccation, locusts infected with *Malpighamoeba* would need to enhance water reabsorption to counterbalance the increased fluid secretion rate of the Malpighian tubules. So, infected locusts may be subjected to the increased energy costs of reabsorbing the excess of water and ions transported into the lumen tubule. Thus, *Malpighamoeba* infection may be highly debilitating for desert locusts in their natural environment, imposing stresses in terms of water, ions, and energy even if they can avoid ingesting plant toxins.

*Malpighamoeba* infections are not limited to locusts and grasshoppers, occurring in other insects including honeybees^11^. Although the exact details of *Malpighamoeba* infection likely depend upon the specific host and pathogen species combination, honeybee Malpighian tubules infected with *Malpighamoeba mellificae* show striking similarities to locust tubules infected with *Malpighamoeba locustae.* In both cases, the lumen of the tubules is swollen and packed with cysts, the epithelium thins, and the brush border of the epithelial cells is destroyed^9,11,12,14^. Such similarities in tubule pathology despite the separation of the insect hosts by approximately 380 million years^45^ suggests that *Malpighamoeba* infection may have similar consequences in other insects susceptible to infection. Moreover, infection in honeybees and other insects may similarly compromise detoxification of xenobiotic compounds and increase water and ion loss. Even though xenobiotic compounds may be removed from insect haemolymph by infected tubules, our results show a substantial increase in fluid secretion by infected tubules, which implies greater energy costs for reabsorption in the insect hind gut. Such impacts are highly injurious and can lead to the premature death of the insect^46^.

## Data accessibility

All supporting data can be found in the electronic supplementary information.

## Acknowledgements

We thank Laura Corona for comments on an earlier version of the manuscript, and the two anonymous reviewers for their valuable comments.

## Competing interests

The authors declare no competing or financial interests.

## Author contributions

M.R. and J.E.N. designed the experiments and wrote the paper. M.R. performed the experiments and final data analysis. S.R.O. provided the infected locusts, devised the statistical analyses and contributed to writing the manuscript. All authors contributed to the interpretation of the analyses.

## Funding

M.R. was funded by a University of Sussex School studentship. S.R.O. was funded by research project grant BB/L02389X/1 by the BBSRC. J.E.N. was funded by research project grant BB/R005036/1 by the BBSRC.

